# Visual processing mode switching regulated by VIP cells

**DOI:** 10.1101/084632

**Authors:** Jung Hoon Lee, Stefan Mihalas

## Abstract

The responses of neurons in mouse primary visual cortex (V1) to visual stimuli depend on behavioral states. Specifically, surround suppression is reduced during locomotion. Although locomotion-induced vasoactive intestinal polypeptide positive (VIP) interneuron depolarization can account for the reduction of surround suppression, the functions of VIP cell depolarization are not fully understood. Here we utilize a firing rate model and a computational model to elucidate the potential functions of VIP cell depolarization during locomotion. Our analyses suggest 1) that surround suppression sharpens the visual responses in V1 to a stationary scene, 2) that depolarized VIP cells enhance V1 responses to moving objects by reducing self-induced surround suppression and 3) that during locomotion V1 neuron responses to some features of the moving objects can be selectively enhanced. Thus, VIP cells regulate surround suppression to allow pyramidal neurons to optimally encode visual information independent of behavioral state.

## Introduction

Visual perception, an internal model of external environment, does not merely reflect exogenous stimuli. Instead, it depends on various endogenous contexts. Consider a number of striking studies in mouse visual cortex that suggest that contextual information originating from other cortical areas modulates primary visual cortex (V1) neuron responses by way of vasoactive intestinal polypeptide positive (VIP) interneurons^1–4^. For instance, the cingulate area (Cg), which modulates the gain of V1 neurons, induces excitatory postsynaptic potentials in VIP cells^1^ as it occurs during locomotion^2^. Thus, it is imperative to comprehend how VIP cells contribute to contextual modulation of V1 neuron responses.

VIP cells, one of the major inhibitory cell types in neocortex^5,6^, are commonly found in superficial layers^7^. They preferentially inhibit somatostatin positive (SST) cells that mediate surround suppression^8,9^. That is, depolarized VIP cells disinhibit pyramidal (Pyr) cells by lowering surrounding suppression. This disinhibition, in fact, accounts for the reduction of surround suppression during locomotion^2,10^. However, it remains unclear why surround suppression is reduced during locomotion. When an animal moves forward, the entire scene, including all objects, appears to move backward (optical flow). When the image of an object moves over the retina, it stimulates multiple receptive fields. As the center of one receptive field constitutes the surround of nearby receptive fields, this motion can induce surround suppression among these cells, a phenomenon we refer to as *self-induced surround suppression*. Thus, the responses of visual-selective neurons to object motion will depend on the strength of *self-induced surround suppression*.

During locomotion, surround suppression in V1 can become too strong for V1 neurons to respond properly to visual stimuli, as all objects are in relative motion. Thus, we hypothesize that VIP cells are depolarized to reduce such surround suppression which may be undesirable during locomotion. To address this hypothesis, we utilize a simple neuronal circuit model of V1, in which the three major inhibitory cell types, parvalbumin (PV), SST and VIP positive inhibitory interneurons, interact with one another and with pyramidal (Pyr) cells via cell-type specific connections^8,9^. We estimate the strength of *self-induced surround suppression* in V1 and demonstrate how VIP cell depolarization enhances visual responses during locomotion by suppressing it. Furthermore, our firing rate and computational models predict that V1 neuron responses to behaviorally relevant features are selectively enhanced during locomotion.

## Results

To address our hypothesis, we first use a firing rate model to study the function of surround suppression and investigate how VIP cell depolarization during locomotion modulates visual neuron responses. Then, we use a computational model of V1 to further validate the findings of firing rate model pertinent to the functions of locomotion-induced VIP cell depolarization. The first subsection describes the numerical analyses of the firing rate model, and the second subsection discusses the computational model simulations.

### Firing rate model and 1-dimensional visual scene

The firing rate model considers a 1-dimensional chain of populations which is connected to 1-dimensional retina (Fig. 1a). In each population, the four cell types are connected via cell type specific connections (Fig. 1b and Supplementary Table 1). All cell types receive tonic external background inputs, which controls their excitability. That is, the strengths are dependent on the cell types (Supplementary Table 2) and are independent of the populations. The firing rate of cell types obey the simple dynamics (Equation 1). The gain function of the firing rate model captures the characteristics of the F-I curve of a leaky and integrate fire neuron model (Supplementary Fig. 1a). This gain function is an approximation rather than the exact F-I curve, but it is less computationally intensive than the exact F-I curve and one of the commonly used gain functions^11^. The synaptic inputs (Equation 2) are the products of weights and gating variables that evolve over time (Equation 3).

**Fig. 1:**
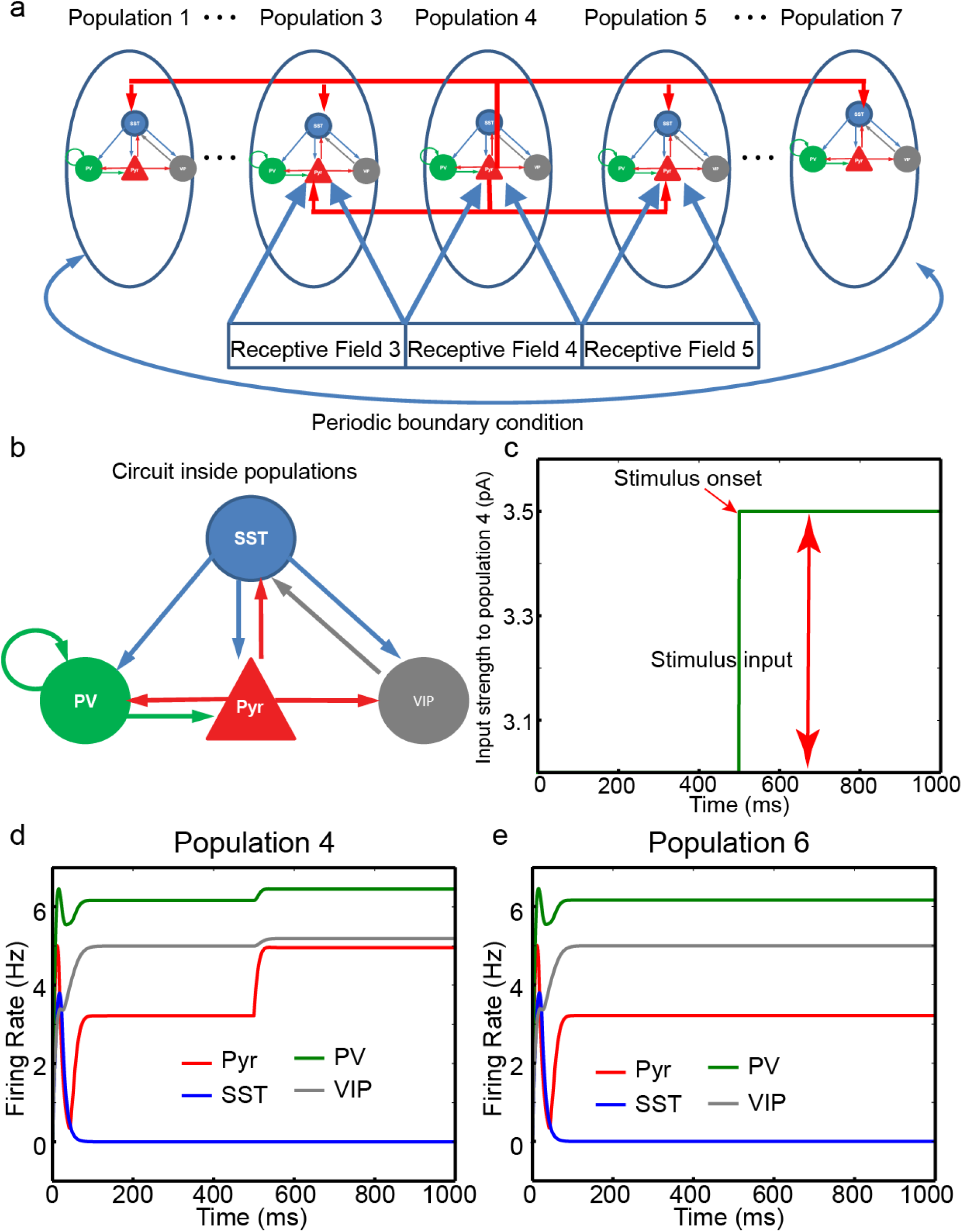
The model structure of firing rate model and its response. **(a)**, 7 populations are implemented in the firing rate model, and each population consists of Pyr, SST, PV and VIP cells. They interact with one another via cell type specific connectivity displayed in **(b)**. Pyr, PV, SST and VIP are shown in red, green, blue and gray, respectively. **(c)**, The time course of input to Pyr cell in population 4. The onset of stimulus presentation is 500 ms marked by the red arrow. **(d)** and **(e)**, Cell type specific activity in populations 4 and 6, respectively, when all inter-population Pyr-SST connections (IPPS) are removed. All other populations, which are not shown explicitly, have identical responses to those of populations 6.

In each population, the four cell types interact with one another, and this “local” circuit in a single population generates rich dynamics^12^. With synaptic events evolving over time (Equation 3), the decay time constants can also modulate the behaviors of this local circuit. To better understand their effects on Pyr cell responses, we performed bifurcation analyses with XPPAUT^13^. Interestingly, we note that the decay time constants of connections from SST to VIP cells (SST-VIP) and from VIP to SST cells (VIP-SST) modulate the Pyr cell response and its stability. As the decay time constant of SST-VIP connection increases, Pyr cell response decreases (Supplementary Fig. 1b). In contrast, as the decay time constant of VIP-SST connections increases, Pyr cell response increases (Supplementary Fig. 1c). This local circuit is stable (red line) in the vicinity of default values of decay time constants, but otherwise they become unstable (black line). At the transition of stability, Pyr cell responses become oscillatory; this oscillatory behavior is induced when SST cell activity is enhanced (see insets of Supplementary Figs. 1b and c).

In the model, 7 populations interact with one another via short-range Pyr-Pyr and long-range Pyr-SST connections known to mediate surround suppression^8^. As seen in Fig. 1a, we establish reciprocal inter-population Pyr-Pyr connections between the two nearest neighboring populations only and inter-population Pyr-SST connections among all populations, as in the earlier computational models^14,15^; in those earlier models, only generic inhibitory cells were considered. The periodic boundary condition is used to ensure all populations are identical in terms of inter-population synaptic inputs. For simplicity, we assume each population is connected to non-overlapping spatial receptive field (RF) which maps onto 1-dimensional visual scene. (Fig. 1a).

### Surround suppression can sharpen responses to a static visual scene

To examine the effects of surround suppression on visual responses, we investigate how it modulates neural responses to an object covering the RF of population 4. This visual object is simulated by providing an additional input (0.5 pA) to Pyr cells in population 4, and it is turned on at 500 ms (Fig. 1c). Due to the background input to Pyr cells, Pyr cells in population 4 receive 3.5 pA input, whereas all other Pyr cells receive 3.0 pA input. In this numerical analysis, we gradually increase the strength of inter-population Pyr-SST connections (IPPS) from 0 pA. When the IPPS strength is set to 0, the firing rates of all cell types reach their steady states after transient responses lasting ~100 ms (Figs. 1d and e). As expected, Pyr cell activity in population 4 is enhanced at 500 ms, and its elevation is bigger than that of the input, as the recurrent connections among Pyr cells provide positive feedback inputs (Supplementary Table 1). All other populations do not show conspicuous changes in response to the stimulus input as shown in Fig. 1e.

More importantly, the IPPS strength has a strong impact on visual responses in the model. When its strength is increased to 15 pA, Pyr cell activity in population 4 is only transiently increased by the stimulus input and then reduced even below its baseline 200-500 ms (Fig. 2a). SST cell activity in population 4 is also enhanced by the stimulus input (Fig. 3a), but this enhancement is not observed in other populations (Fig. 2b); all populations except population 4 show identical responses. In population 4, at the onset of the stimulus input, Pyr cell activity is enhanced, increasing the synaptic excitation to SST cells. Although population 4 of Pyr cells send excitation onto all SST cells, it drives population 4 of SST cells most strongly (see Supplementary Table 1). With this strong local drive within population 4, SST cell activity is elevated (Fig. 2a), but in all other populations except population 4, SST cell activity remains unmodulated by the stimulus input (Fig. 2b); that is, IPPS is not strong enough to excite SST cells in other populations but population 4. As SST cell activity increases, the firing rates of all other cell types decrease. Even though Pyr cell activity is below its baseline, the elevated SST cell activity is sustained because of the reduction of inhibition from VIP to SST cells (Fig. 2a).

**Fig. 2:**
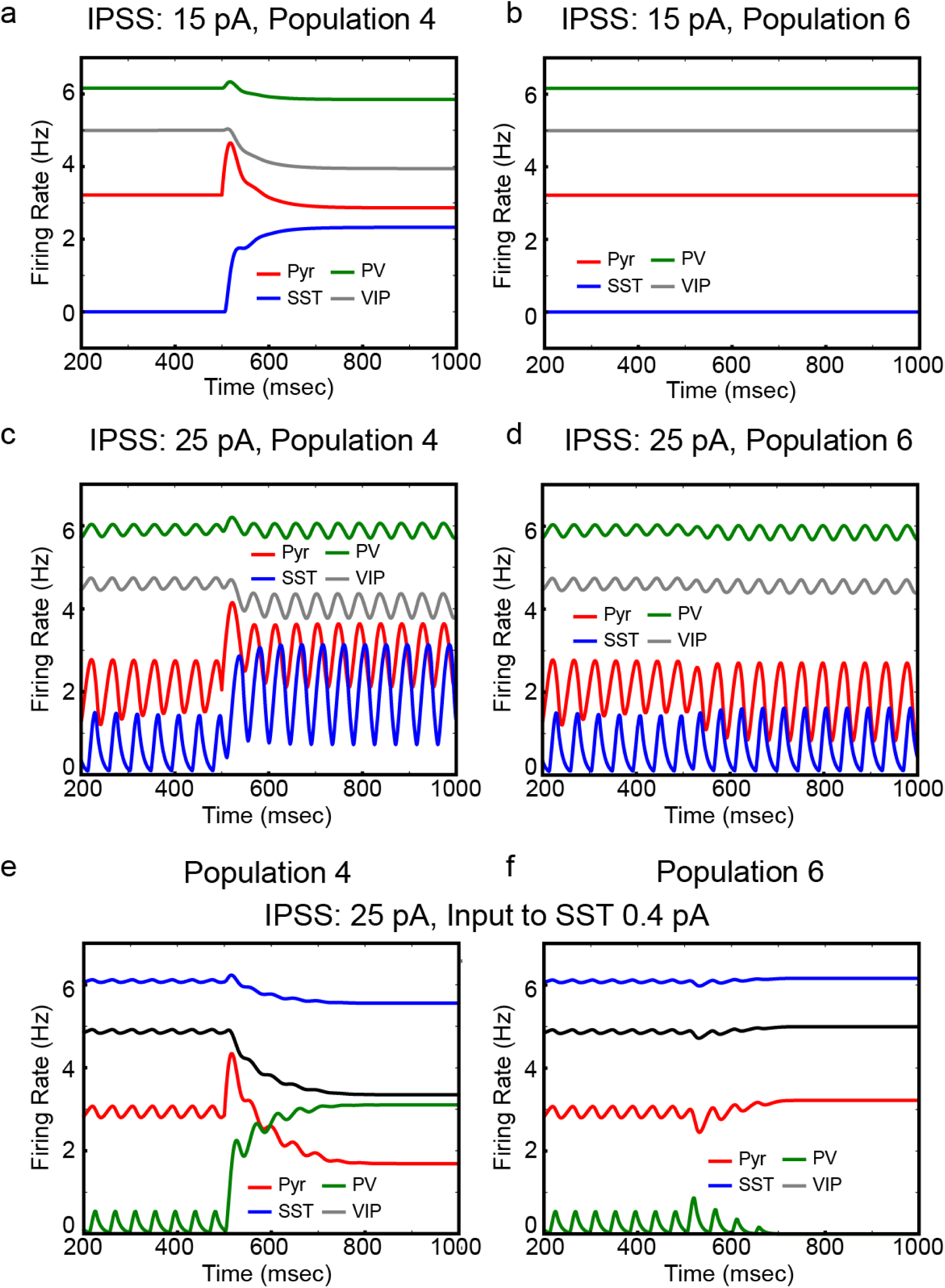
The effects of surround suppression on model responses. **(a)** and **(b)**, Cell-type specificactivity in populations 4 and 6, respectively, when the strength of IPPS is 15 pA. **(c)** and **(d)**, The same but with enhanced IPPS strength (25 pA). **(e)** and **(f)**, The same when the background input to SST cells and IPSS strength are 0.4 and 25 pA, respectively.

**Fig. 3:**
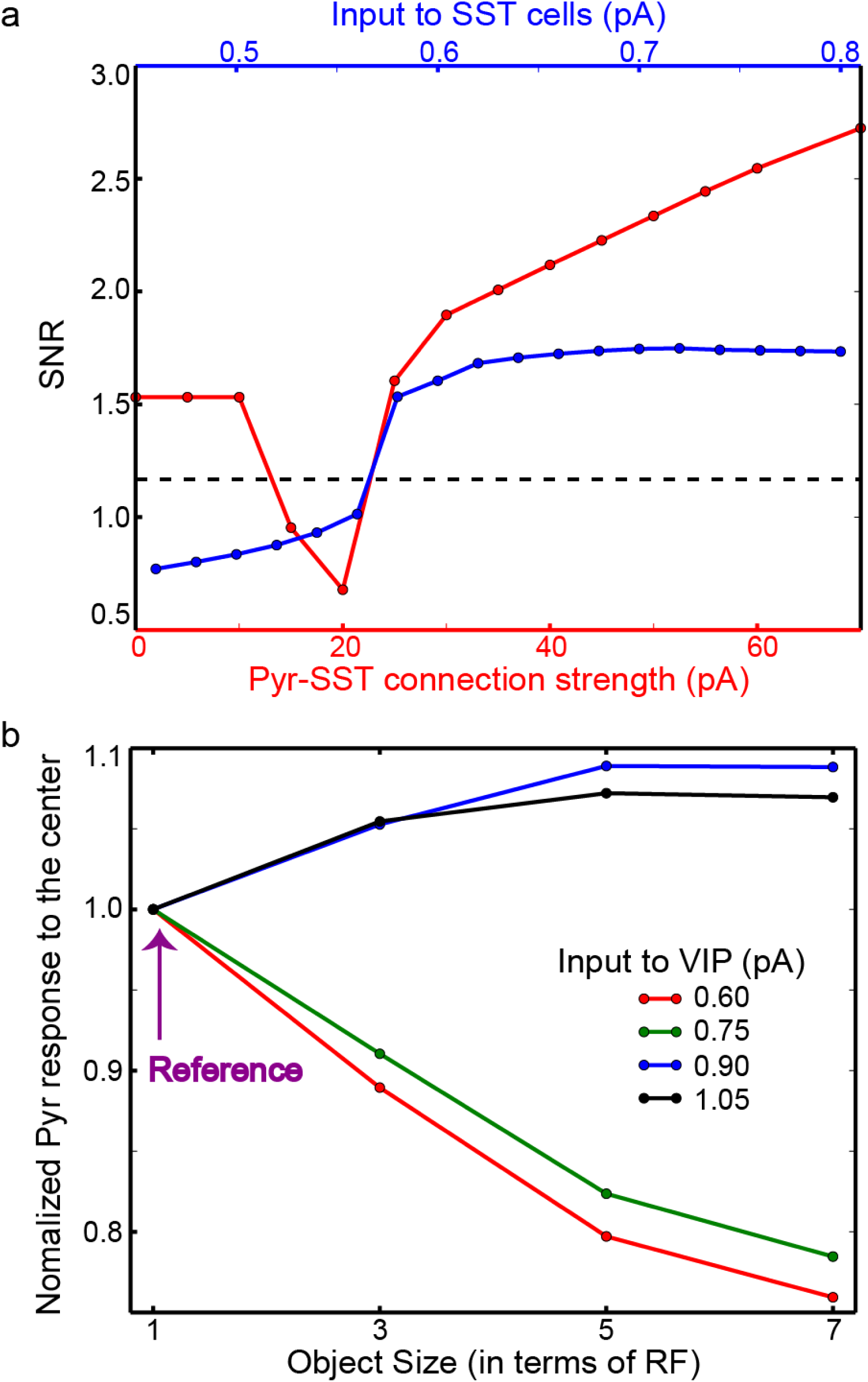
The functions of surround suppression and its modulation. **(a)**, Pyr cell responses inpopulation 4, compared with other Pyr cell responses elicited by background input only. SNR is the normalized population 4 responses to the mean activity of all other populations. The dependency of SNR on the input to SST cells and the IPPS strength are shown in blue and red, respectively. The dashed line represents the SNR of inputs to Pyr cells. **(b)**, The modulation of surround suppression via VIP cell depolarization. *y*-axis represents Pyr cell responses to the center of the object depending on its size. For a fixed input to VIP cells, we set the Pyr cell responses to the narrow object, whose width is 1-RF long, as reference values, which we use to normalize Pyr cell responses. The background inputs to VIP cells are 0.6, 0.75, 0.9 and 1.05 pA, which are shown in red, green, blue and black, respectively.

When the IPPS strength is further enhanced to 25 pA, the model shows strikingly different behaviors. First, the responses become oscillatory (Figs. 2c and d), which reflects the intense interactions among populations. The frequency of this oscillation is ~22 Hz (Supplementary Fig. 2a), and this oscillatory behavior is abolished when we hyperpolarize SST cells by introducing inhibitory currents. (Supplementary Figs. 2b and c); the inhibitory currents are introduced to SST cells between 700 and 800 ms, which are marked with a black arrow. Thus, this oscillation is generated by the interplay between SST and Pyr cells, which is consistent with our bifurcation analysis (Supplementary Fig. 1b and c) and the earlier experimental/computational study^16^. Second, as seen in Fig. 2c, Pyr cell activity in population 4 is sustained during the stimulus period (500-1000 ms), and we note a slight decline in Pyr cell activity and a slight surge in SST cell activity in all other populations (see Fig. 2d for an example). As surround suppression is mediated by SST cells, the background input to SST cells can also modulate surround suppression. Its effects are indeed consistent with those of IPPS. When the background input to SST cells is reduced to 0.4 pA, Pyr cell activity in population 4 is reduced during the stimulus period (Fig. 2e), as Fig. 2a shows Pyr cell activity with the weaker IPPS. For comparison, we display population 6 responses in Fig. 2f.

Interestingly, Pyr cell responses induced by the sensory input (i.e., Pyr cell response in population 4) can be either stronger or weaker than those in other populations depending on the IPPS strength. With IPPS=25 pA (Figs. 2c and d), Pyr cell activity during the stimulus period (500-1000 ms) is much stronger in population 4 than in other populations. In contrast, with IPPS=15 pA (Figs. 2a and b), Pyr cell activity in population 4 is weaker than that in other populations. These results suggest that the stimulus input to population 4 exerts inhibition to other populations via SST cells only when IPPS is strong enough. To address this further, we quantify how strongly the stimulus input drives Pyr cells in population 4, compared with others. Specifically, we calculate the signal-to-noise ratio (SNR) by normalizing Pyr cell activity in population 4 to the mean value of Pyr cell activity in other 6 populations; that is, we estimate the stimulus-evoked Pyr cell activity relative to the background input-driven Pyr cell activity.

The blue and red lines in Fig. 3a show the dependency of SNR on the background input to SST cells and the strength of IPPS, respectively. When IPPS strength is less than 10 pA, IPPS has little impact on model responses. However, when the strength of IPPS is 15 or 20 pA, Pyr cell responses in population 4 are weaker than those in other populations. This is due to the selectively enhanced SST cell activity (Figs. 2a and b); that is, in these regimes, the feedback inhibition from SST to Pyr cells is prominent in population 4 only, and thus SNR is smaller than 1. When IPPS is further strengthened, SST cells in other populations start firing and mediate lateral inhibition (i.e., surround suppression), and Pyr cell responses in population 4 are stronger than those in other populations (Figs. 2c and d); that is, the visual response are sharper. As the strength of IPPS grows, SNR increases (Fig. 3a). We also normalize the stimulus-evoked response (500-1000 ms) to the baseline-period activity (200-500 ms) for each population. Specifically, we calculate the mean Pyr cell activity in both periods and estimate the relative changes (Equation 4). As seen in Supplementary Figs. 3a and b, the stimulus evoked activity relative to the baseline activity is consistently modulated in the way SNR is modulated (Fig. 3a). These results indicate that surround suppression mediated by SST cells makes visual responses to the object sharper only when IPPS is strong enough.

Next, we study how surround suppression is dependent on the decay time constants of connections from SST to Pyr cells (SST-Pyr) and from SST to VIP cells (SST-VIP). SNR values in Supplementary Fig. 3c show that surround suppression become more effective when SST-Pyr inhibition is prolonged. When the decay time constant of SST-Pyr inhibition is shorter than 6.5 ms and longer than 5 ms, SST-Pyr inhibition becomes effective only in population 4, in which SST cells are sufficiently active. That is, Pyr cells in population 4 receive additional inhibition, making SNR below 1 in this regime. When SST-VIP inhibition is prolonged, SST cell activity increases (Supplementary Fig.1b), and thus the inhibition of SST impinging onto Pyr cells is enhanced. This enhanced inhibition onto Pyr cells suppresses stimulus evoked responses, which accounts for the negative correlations between SNR and the decay time constant of SST-VIP inhibition.

We also examine whether VIP cell depolarization could reduce surround suppression. To do so, we measure how the firing rate model of Pyr cells in population 4 is modulated by the size of visual object. In the four experiments, 1 RF-, 3 RF-, 5 RF-and 7 RF-long objects are presented, respectively. In each experiment, the center of the object always stimulates population 4, and Pyr cell responses in population 4 are measured between 500-1000 ms. That is, we simulate the standard estimation of surround suppression strength. As seen in Fig. 3b, in the model, VIP cell depolarization can reduce surround suppression. When the input to VIP cells is weak, Pyr cell response to the center (Pyr cell response in population 4) declines, as the size of the object grows. In contrast, Pyr cell response to the center becomes stronger when the input to VIP cells is increased to 0.9 pA and higher. That is, when VIP cells are depolarized, surround facilitation emerges instead of surround suppression.

### VIP cell depolarization can enhance visual responses during locomotion

Next, we ask: how does VIP cell depolarization modulate visual neuron responses during locomotion? When a mouse is running, we expect some objects to move towards the mouse and others to move away. Below, we examine both possibilities.

First, we consider an object moving away. In this condition, a 3 RF-long object is assumed to move to the right (Fig. 4a), and we examine Pyr cells’ response to it depending on the input to VIP cells. At every 50 ms we update the object’s location by 25% of receptive field size. The stimulus input is proportional to the area of receptive field covered by the object. That is, population 1 receives the full sensory input (0.5 pA) during 300-350 ms, but this input decreases gradually by 25% at every 50 ms (Figs. 4b). In contrast, population 4 receives gradually increasing sensory inputs, as the object is approaching the RF of population 4. At 500 ms, population 4 receives the full sensory input. We remove the object from the scene at 550 ms. As a control experiment, we examine Pyr cell responses to the moving object without surround suppression (i.e., no IPPS). As seen in Fig. 4c, Pyr cell responses faithfully reflect the stimulus input. To assess the effects of surround suppression, we restore surround suppression and estimate Pyr cell responses depending on VIP cell depolarizations (Figs. 4d and e). In those figures, the Pyr cell responses are normalized to the maximum response during simulations and are indicated in color; the red represents the maximum response. The surround suppression globally reduces Pyr cell responses (Fig. 4d). Specifically, Pyr cells responses are prominent only between 300-350 ms yet decrease afterwards, supporting our hypothesis that self-induced surround suppression reduces Pyr cells’ sensitivity to moving objects. When the input to VIP cells is increased to 1.2 pA, population 4 responses are stronger than other populations during the stimulus period of the entire movement (Fig. 4e). That is, VIP cell depolarization almost exclusively enhances responses to RF 4, toward which the object moves.

**Fig. 4:**
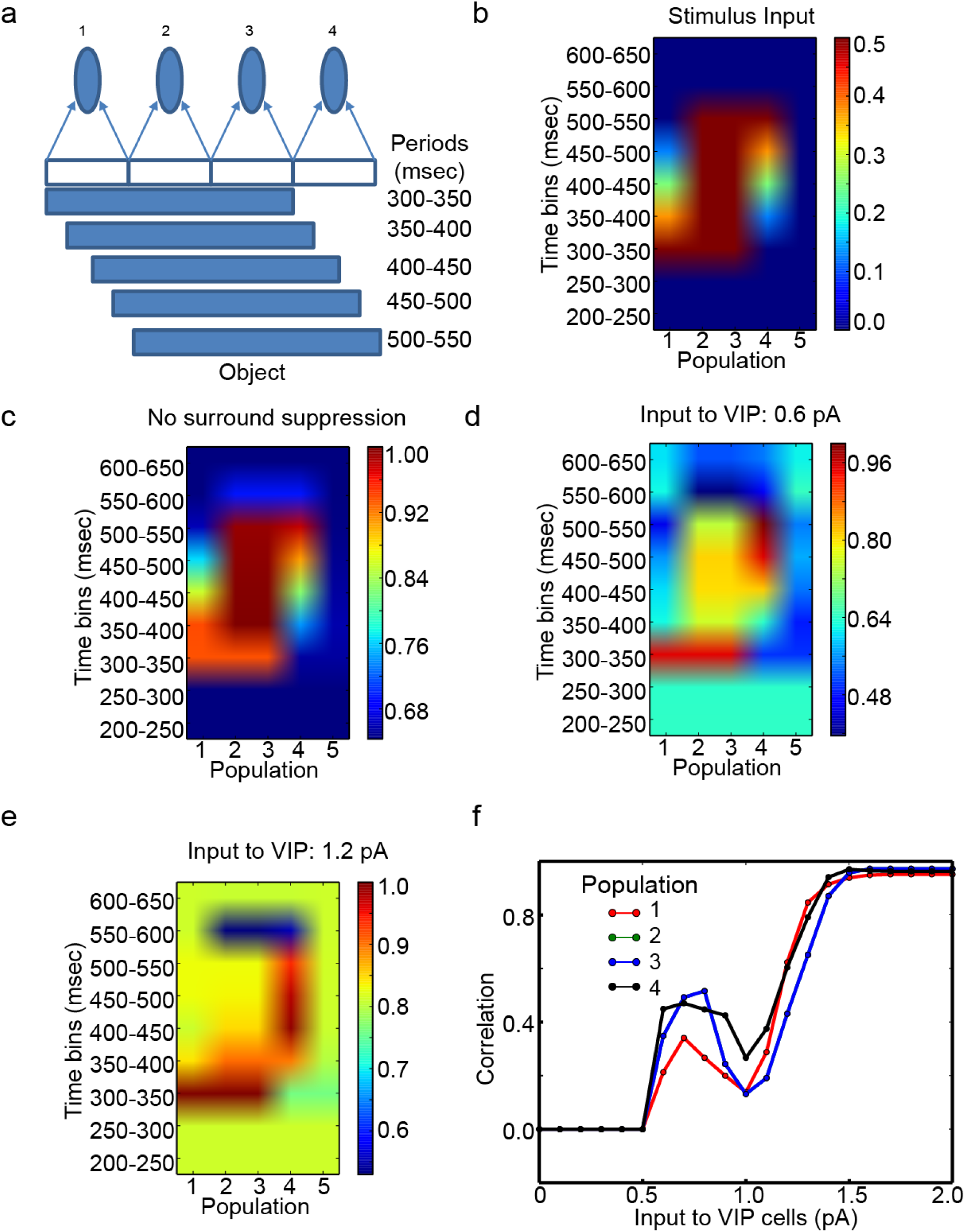
The effects of VIP cell depolarization on model responses to the object in motion. **(a)**, Method by which we simulate the object moving over time. **(b)**, Stimulus input introduced to the populations in color codes. The red indicates the maximum stimulus input 0.5 pA. The *x*- and *y*-axes represent the identity of population and time bins. **(c)**, The normalized Pyr cell responses to the stimulus input shown in **(b)** when all inter-population Pyr-SST cell connections are removed. **(d)** and **(e)**, The normalized Pyr cell responses with surround suppression depending on the input to VIP cells. In (c)-(e), Pyr cell responses are divided by the maximum Pyr cell responses (over all populations) during simulations. The strength of the input to VIP cells is shown above each panel. **(f)**, Population specific correlations between stimulus inputs and the Pyr cell responses. As the population 2 and 3 show identical responses, only population 3 is visible in (f).

To better understand the effects of VIP cell depolarization on visual responses, we quantify how reliably Pyr cell outputs reflect the stimulus inputs by calculating Pearson’s correlation between inputs to Pyr cells and their outputs in each population. If Pyr cell outputs depend on the inputs completely, the correlation should be 1. There are three different regimes (Fig. 4f); populations 2 and 3 show identical responses, and thus population 2 is not visible in the figure. In the first regime, in which the input to VIP cells is lower than 0.6 pA, Pyr cells are quiescent. While their firing rates and the covariance between the inputs to and outputs from Pyr cells are both below 10^−7^, we observe noticeable correlations, which are ~ −0.1, in this regime. To avoid any possible artifacts from this tiny yet non-zero Pyr cell activity, we display the covariance instead of the correlation (Fig. 4f) when the covariance is below 10^−7^. In the second regime, in which the input to VIP cells is between 0.6 and 1.1 pA, the population output becomes less dependent on the input, as the input to VIP cells increases. As populations 1 and 4 receive the same amount of total inputs during the simulation period, we can directly compare the correlation between them. As seen in Fig. 4f, population 4 output reflects its input more faithfully than population 1 when the input to VIP cells is between 0.6 and 1.1 pA. Additionally, the correlation of population 4 is the highest, when the input to VIP cells is 0.9 or 1.0 pA. In the third regime, in which the input to VIP cells is bigger than 1.5 pA, all correlations increase and converge to 1. The most intriguing observation is that the correlations are dissimilar among populations in the second parameter regime, suggesting that VIP cell depolarization can selectively enhance visual responses rather than uniformly.

Second, we consider an object approaching the mouse. The approaching object is simulated by increasing its size over time (Fig. 5a). Specifically, the number of populations stimulated by this object increases over time. Population 3 receives the stimulus input (0.5 pA) between 300 and 600 ms, populations 2 and 4 receive it between 400-600 ms, and populations 1 and 5 receive it between 500-600 ms. As seen in Fig. 5b, we note that Pyr cell activity depends on the input to VIP cells; populations 1 and 5 show identical responses with each other, and populations 2 and 4 also show identical responses, and thus populations 1 and 2 are not visible in Fig. 5b. When the input to VIP cell is low (0.6 pA), Pyr cell activity in population 3 is elevated at 300 ms, which reduces over time (Fig. 5B), even though population 3 receives constant stimulus inputs between 300 and 600 ms. This reduction disappears when the input to VIP cells is increased to 1.8 pA (Fig. 5b). Interestingly, the reduction seems more pronounced when the input to VIP cells is at an intermediate level (1.2 pA). We again calculate the correlations between inputs to and outputs from Pyr cells depending on the input to VIP cells. As in Fig. 4, we show the covariance instead of the correlation when it is smaller than 10^−7^. As seen in Fig. 5c, the correlations are modulated by the input to VIP cells. The correlations of populations 1 and 5 almost monotonically increases, as the input to VIP cells increases. In contrast, the correlations of other populations increase in the beginning until the input to VIP reaches a certain threshold value, and they start decreasing (Fig. 5c). When the input is close to 2 pA, the correlations of all populations approach 1.0.

**Fig. 5:**
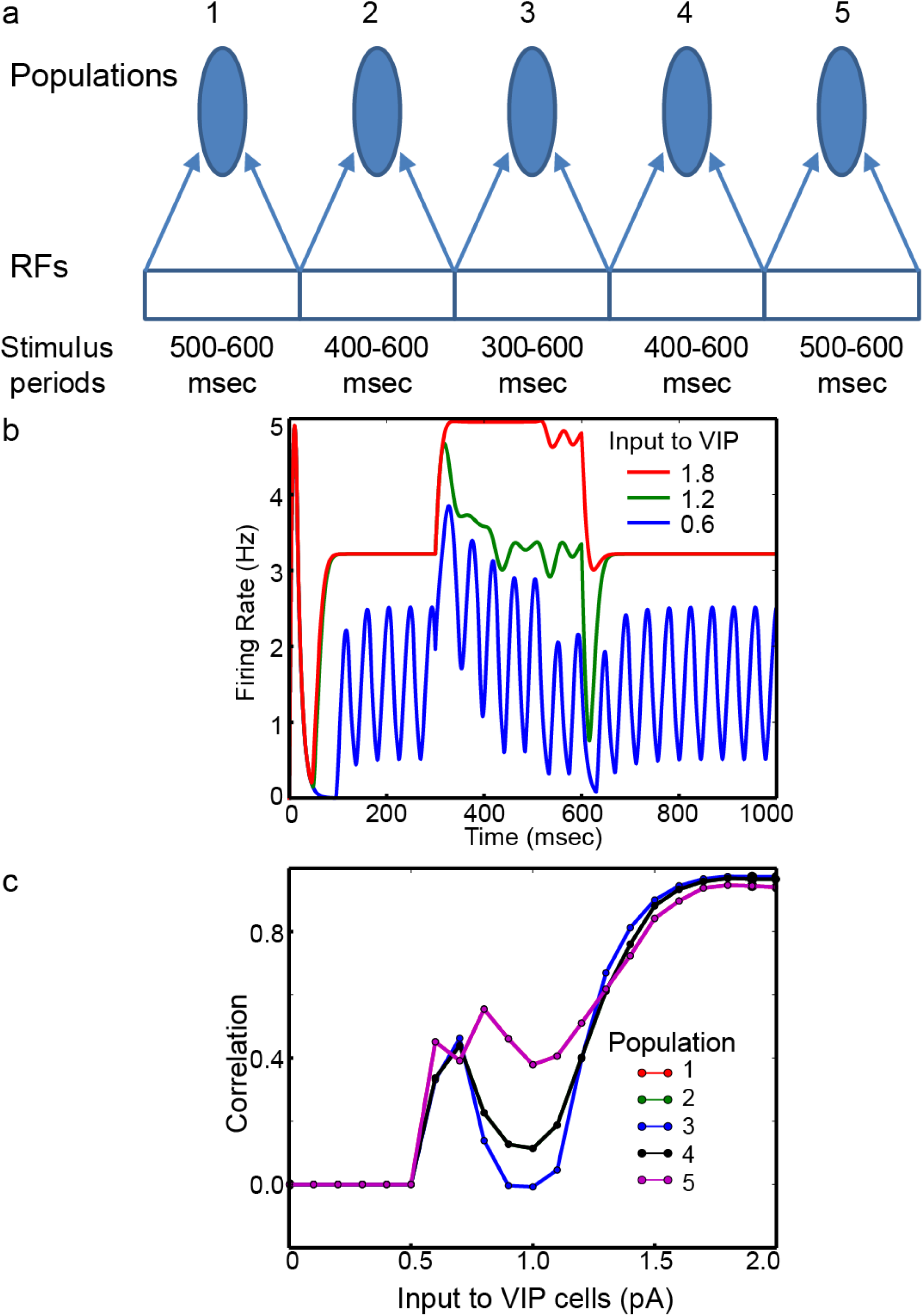
The effects of VIP cell depolarization on responses to the object growing in size. **(a)**, Method by which we simulate the object growing in size. **(b)**, Pyr cell activity in population 4 between low (0.6 pA) and high (1.8 pA) input to VIP cells. For comparison, Pyr cell activity with the intermediate input (1.2 pA) to VIP cells is also displayed. **(c)**, Dependency of population specific correlations between the stimulus inputs and Pyr cell responses on the input to VIP cells. Populations are distinguished using different colors. As populations 2 and 4 (1 and 5) show identical responses, only three lines are visible in (c)

In brief, we note 1) that surround suppression leads to sharper visual responses to stationary visual scene, 2) that VIP cell depolarization may help V1 cells respond to objects in motion and 3) that the benefit of VIP cell depolarization may not be homogenous. Instead, VIP cell depolarization selectively enhances visual responses to some features (Figs. 4f and 5c) when the input to VIP cells is intermediate.

### Computational model with 2-D visual scene

The numerical analyses of the firing rate model indicate that locomotion-induced VIP cell depolarization effect is feature-specific. However, we cannot exclude the possibility that these results are artifacts attributable to either 1) the firing rate model that provides a qualitative approximation of neural dynamics rather than exact description, or 2) the abstract 1-dimensional visual scene. Thus, to further validate these findings, we use a computational model of V1 responding to a more realistic 2-dimensional visual scene. The computational model used here is an extension of our earlier model^12^, in which, PV, SST and VIP cells in the superficial layers of 13 columns interact with one another and with Pyr cells via cell-type specific connections within and across columns. The earlier model^12^ also includes long-range and short-range inhibitions across columns mediated by SST and PV cells, respectively. Maintaining the inhibitory cell types and cell-type connectivity of the earlier model, we extend it into a 2-dimensional array of 192 cortical columns, each of which has ~2000 cells, as shown in Fig. 6a, to test V1 responses to a more realistic 2-dimensional visual scene (Methods).

**Fig. 6:**
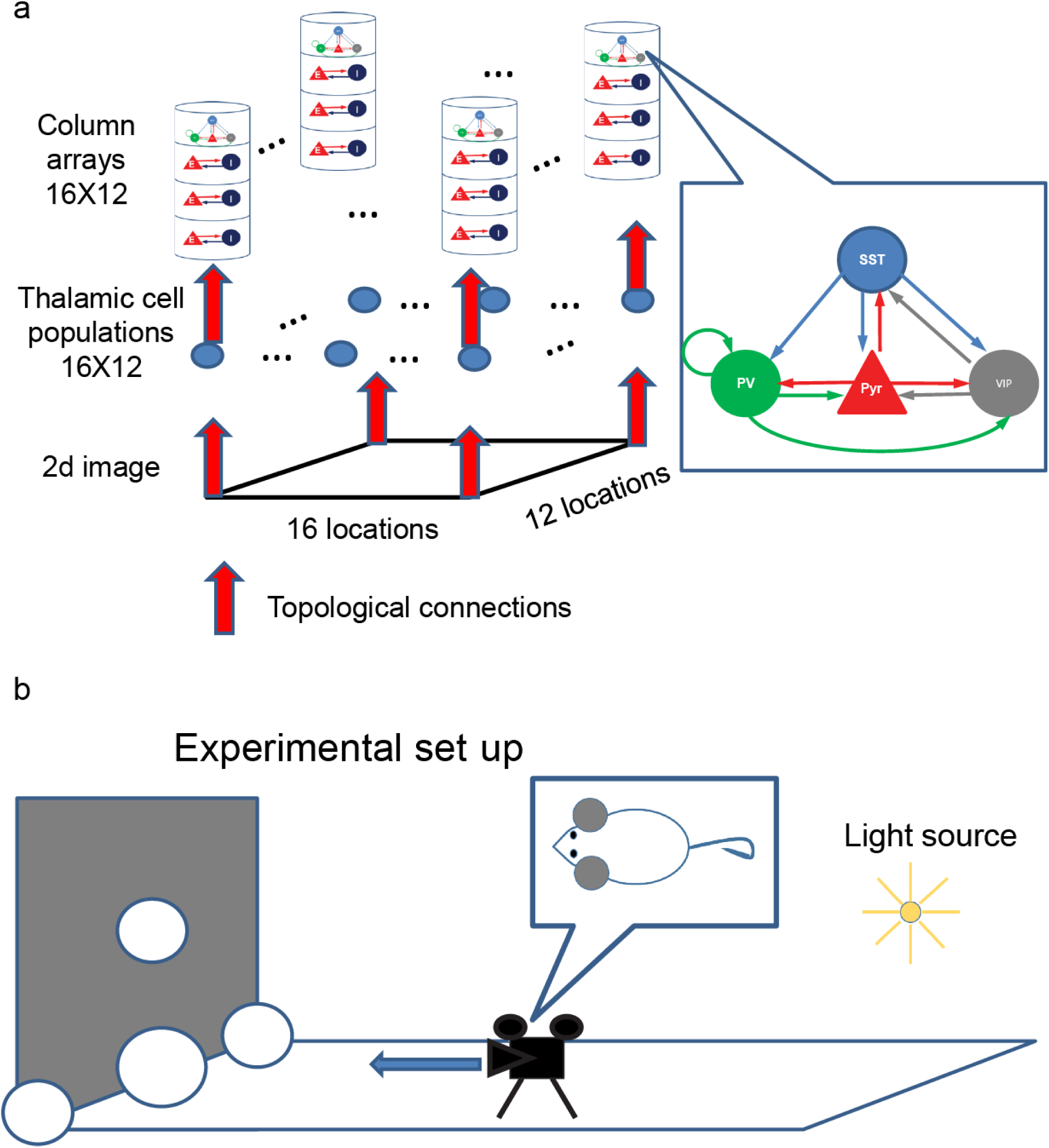
The model structure of computational model of V1. **(a)**, Structure of the computationalmodel. In the computational model, we consider more connections among the four cell types^12^ than those used in the firing rate model. The model consists of 192 columns distributed over a 16-by-12 grid. The receptive field of each column covers non-overlapping image patches consisting of 40-by-40 pixels; the rendered images consist of 640-by-480 pixels. All inter-columnar connections are established using a periodic boundary condition; see Supplementary Table 3. **(b)**, Virtual experiment-setup.

In this study we consider a simple experimental set up, in which a mouse faces a fronto-parallel plane and translates at a constant speed toward this plane (Fig. 6b). This gives rise to a linear flow field with a central focus-of-expansion. We simulate this setup using an image of five spheres (Fig. 6b) and use POV-Ray^17^ to render how they appear to the mouse when the animal runs toward the screen. Images are rendered for 1 sec at 20 frames/sec. In order to focus on the essential nature of the cortical circuit processing, we use highly simplified non-temporal receptive field for both lateral geniculate nucleus (LGN) and cortical neurons. That is, each 50 ms-long frame is spilt into 16-by-12 non-overlapping spatial patches which are mapped topographically onto a population of 100 LGN cells per patch (Fig. 6a). LGN cells in turn send Poisson spike trains to cortical columns in a topographic manner; the connection probabilities for such thalamo-cortical connections are taken from an earlier model^15^. The firing rates of each LGN population of 100 cells are proportional to the sum of light intensities in the corresponding image patch (Equation 5 in Methods) and are updated every 50 ms. We run simulations for 1 sec and record spikes from 10% of layer 2/3 pyramidal cells.

### Depolarized VIP cells modulate V1 neurons to the moving objects in inhomogeneous ways

We assume that a mouse moves at a constant speed toward the central sphere to which is referred as target sphere hereafter (Supplementary Fig. 4); the target sphere is 50% bigger than others. That is, the target sphere grows in terms of size, and all others move outward (the left column of Supplementary Fig. 4). LGN outputs faithfully reflect the location of the spheres in motion as shown in the right column of Supplementary Fig. 4. Locomotion-induced VIP cell depolarization is simulated by increasing the background external inputs from 16 Hz to 20 Hz carried by a single external fiber (Supplementary Table 3). Figure 7 compares the responses averaged over 10 independent simulations between high and low VIP cell depolarization conditions. During the early periods, when all five spheres are presented within the visual field, the column responses to the non-target spheres are sharper in the high depolarization condition (Figs. 7a and b). During the later periods, when the target sphere dominates the visual field, we note strikingly different responses to the target sphere between the high and low depolarization conditions (Figs. 7c and d). The responses of columns connected to the target sphere’s edge are stronger than those connected to the target sphere’s surface in the high depolarization condition. In contrast, the responses to the center are stronger than those to the edges in the low depolarization condition. That is, locomotion-induced VIP cell depolarization suppresses V1 neurons responding to the surface of target sphere, which is consistent with the numerical analysis in Fig. 5. We also display the spikes generated by Pyr, PV, SST and VIP cells in response to the two image patches illustrated in Supplementary Fig. 5a. The left and right columns in Supplemental Fig. 5 show the spikes in the low and high depolarization conditions, respectively. Even in the high depolarization condition, VIP cells are active only when they are responding to the patch 2.

**Fig. 7:**
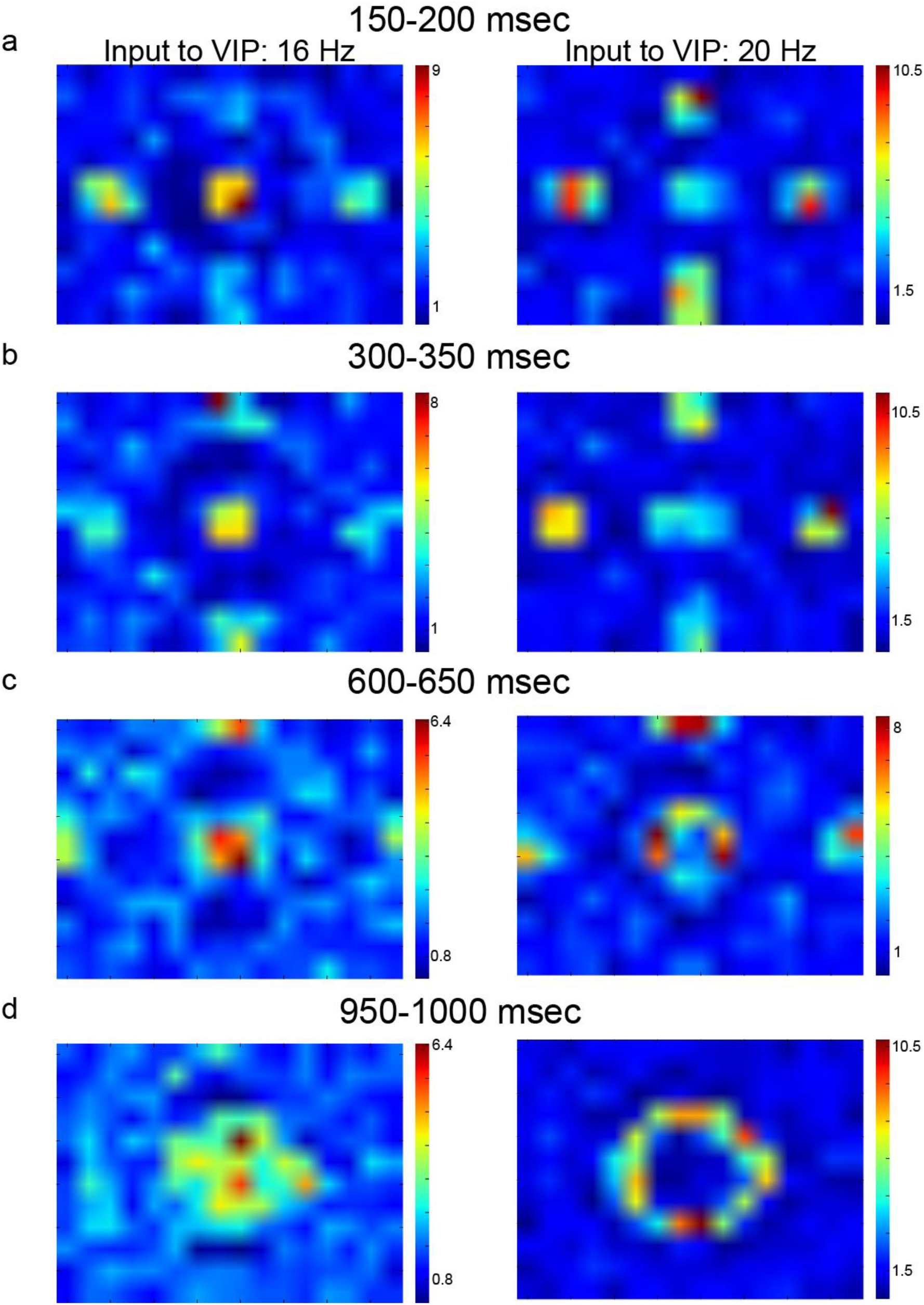
Column responses in the computational model. **(a)-(d)**, Column responses averagedover 10 independent simulations in four different 50 ms time windows, respectively. The left and right columns show them with low and high external inputs to VIP cells. The color bars show the firing rate of Pyr cells in Hz.

To quantify how reliably V1 neurons respond to LGN outputs, we again calculate the correlation between LGN outputs and column responses (recorded spikes from 10 % layer 2/3 Pyr cells); see Methods. As images to LGN cells are updated at every 50 ms, we split column responses to correspond 50 ms-time bins. Then, the correlation is calculated using LGN outputs and column responses in the same time window (Equation 6). The effects of VIP cell depolarization on columns responding to the four spheres moving outward in the visual field seem homogeneous, and thus we do not differentiate them when calculating the correlation. In contrast, the effect of VIP cells on columns responding to the target sphere is clearly distinct from those on columns responding to other spheres. Thus, we split LGN outputs and column responses depending on whether they are induced by the target sphere or not. Specifically, we identify the spatial extent of the target sphere in each frame using thalamic outputs and split the columns and thalamic populations into two distributions (inside-and outside-target distributions). When we analyze the responses induced by the four spheres, we calculate the correlations (COT) between column responses and LGN outputs using the outside-target distribution. For the responses induced by the target sphere, we calculate the correlations (CIT) using the responses in the inside-target distribution. COT is calculated from the first 8 frames, as the four spheres start disappearing from the visual field at 400 ms. In contrast, CIT is calculated from all 20 frames.

CIT would be 1 if Pyr cell activity in the columns connected to the target sphere entirely depends on LGN outputs induced by the target sphere. Otherwise, CIT would be close to 0 if the target sphere cannot drive columns at all. Similarly, COT estimate the capacity of the four spheres to drive V1 neurons via LGN. Fig. 8a shows the estimated correlations (COT and CIT) from 10 independent simulations. As seen in the figure, VIP cell depolarization enhances the COT but reduces CIT, and the induced changes are significant (t-test, p<10^−10^). COT enhancement is consistent with the stronger responses to the four spheres in the high depolarization condition (Fig. 8). CIT reduction is also expected, as the responses to the surface is suppressed in the high depolarization condition. That is, CIT and COT successfully represent the distinct effects between the target and other spheres. Lastly, we estimate how surround suppression modulate CIT and COT to confirm that surround suppression reduction is indeed responsible for the stronger responses to moving objects. In the simulations, we strengthen surround suppression by increasing the connection probability for inter-columnar Pyr-SST connections. If the stronger surround suppression reduces COT, it would support that surround suppression is harmful to visual responses during locomotion. As expected, COT is reduced (Fig. 8b). On the other hand, we note that CIT is increased, suggesting that surface responses are restored when surround suppression is enhanced.

**Fig. 8:**
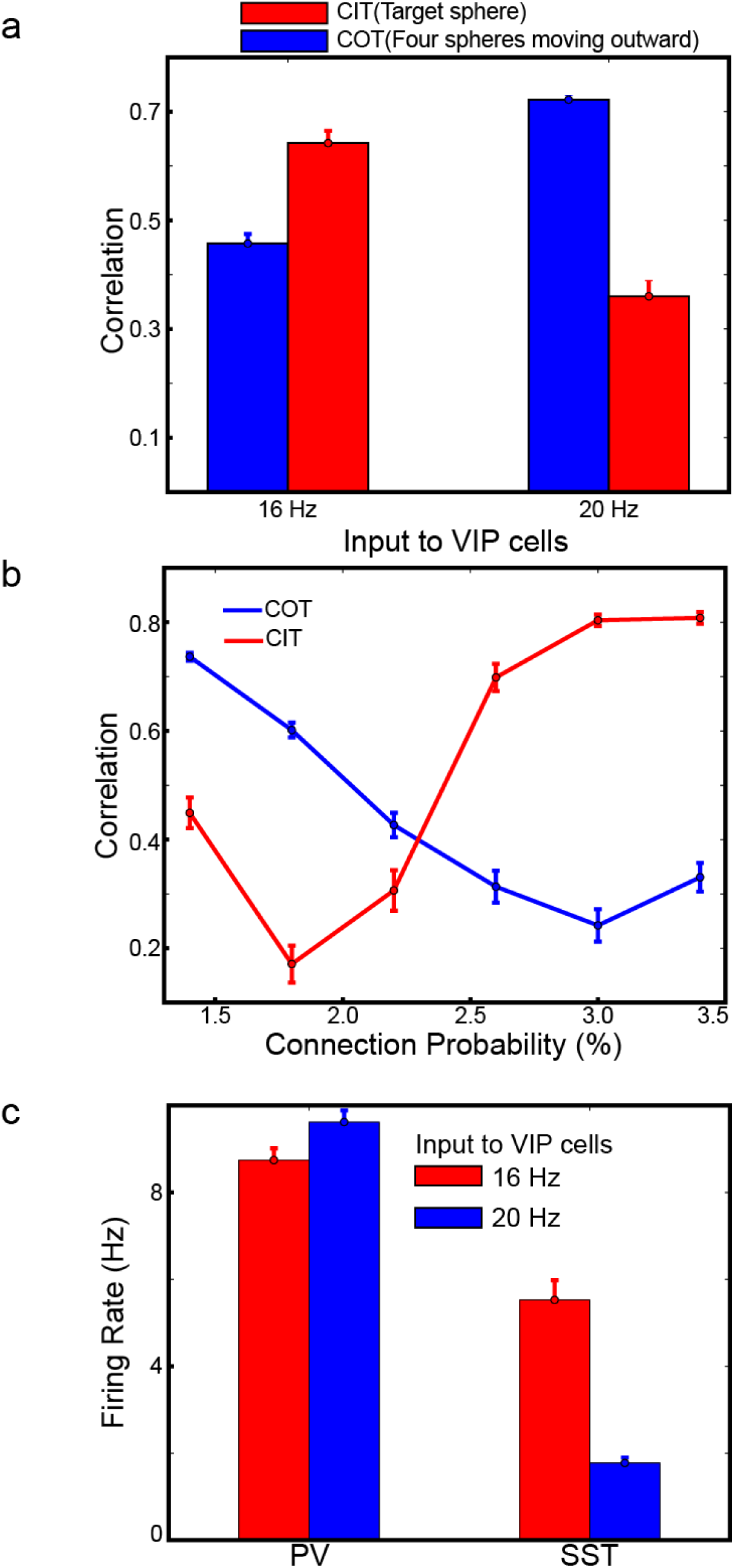
Correlations for the target and other spheres. **(a)**, Correlations between high and lowVIP cell depolarization conditions compared. **(b)**, Dependency of correlations on the surround suppression strength regulated by connection probability for Pyr-SST connections across columns. **(c)**, PV and SST cell activity between high and low VIP cell depolarization conditions. All error bars represent standard errors

### The effects of VIP cell depolarization also modulate SST and PV cell activities

The simulation results exhibit the effects of VIP cell depolarization on Pyr cell responses. Does it modulate other inhibitory cell types? During locomotion, PV cell activity was also reported to be enhanced^2^. In the model, we find a consistent behavior (Fig. 8c) which can be explained by reduced inhibition from SST cells. Interestingly, we note that the enhanced PV cell activity appears necessary to make V1 neurons respond more strongly to the edge of the target sphere than to its surface. When we reduce the background inputs to PV cells, column responses to the surface become stronger, and edge-dominant responses disappear (Supplementary Fig. 6a). We also note that SST cell activity is modulated in a location specific manner despite its reduction in general (Fig. 8c). In the last frame (950 ms-1000 ms), in which only the target sphere exists to dominate the visual field, SST cells responding to the center of the target sphere fire more strongly when VIP cells are depolarized (Supplementary Fig. 6b). Specifically, SST cell activity in the columns connected to the center of the target spheres is increased by ~30%. This can accounts for the recent experimental finding that SST cell activity can also be enhanced during locomotion^18^.

## Discussion

Both firing rate and computational models support our hypothesis that VIP cell depolarization leads to stronger responses to visual objects in relative motion by suppressing self-induced surround suppression during a mouse’s locomotion. The surround suppression promotes sharper responses to stationary visual scene (Fig. 3a). However, it can disrupt visual neuron responses to objects in motion (Figs. 4, 5, 7 and 8b), and VIP cell depolarization is the potential mechanism, by which surround suppression is regulated (Figs. 4, 5 and 7). We note that low-threshold spiking interneurons that express SST are known to burst^16^, which we did not consider in both models. That is, the effects of self-induced surround suppression may be even bigger than those estimated in the models. Below we discuss the implications of our analyses in details.

We emphasize that simulation results of the computational model constrained by experimental data are consistent with the firing rate model responses in the intermediate VIP cell depolarization condition, raising the possibility that visual cortex can indeed work in the regime, in which VIP cell depolarization makes feature-specific enhancement of visual responses. While this suggestion should be examined by future experiments, we propose that such selective enhancement of visual responses may have direct functional advantages. First, the firing rate model suggests that VIP cell depolarization enhances the responses to the RF of population 4 but reduces the responses to the RF of population 1 (Fig. 4). It should be noted that the object moves away from the RF of population 1 and approaches that of population 4. When the object is in motion, its current location may be more crucial than its previous one. The biased enhancement of the RF that receives increasing stimulus inputs makes V1 neurons focus on the current location of the object in motion rather than the previous one.

Second, VIP cell depolarization suppresses the responses to the center of the object growing in size (Fig. 5). The same phenomenon is also observed in the computational model simulations: the responses to the surface of the target sphere growing in size are suppressed, whereas those to the edges are enhanced (Figs. 7 and 8). Importantly, the target sphere in the computational model has a clear behavioral importance as it can collide with a mouse. Thus, the mouse must heed the distance between itself and the sphere. The sphere’s size and its growth rate will be valuable when estimating the distance. It means that the surface of the approaching object could merely be a distraction which can be ignored. In the model, the depolarized VIP cells automatically make V1 neurons ignore the target sphere’s surface (Fig. 7d).

How do V1 neurons become sensitive to some features such as edges/boarders, which represent discontinuity of images? The firing rate model suggests that the interplay between SST and VIP can be a main factor, as selective enhancement of visual responses appears only when the input to VIP cells is neither too strong nor too weak (Figs. 4 and 5). When it is too weak, SST cells are active in all populations, and all Pyr cells become quiescent. When it is too strong, SST cells do not fire, and Pyr cells faithfully respond to stimulus inputs. In the intermediate regime, SST cell activity depends on stimulus inputs, not just to the same population but also to neighboring populations, due to the Pyr-SST connections across populations. Then, it should be noted that SST cells responding to the center of the visual objects will receive the strongest stimulus inputs, as many neighboring populations receive stimulus inputs. That is, Pyr cells responding to the center will be under the strongest inhibition of SST cells, which can account for the suppression of responses to the center of the object (Fig. 5). Also, the location specific modulation of SST cell activity observed in the computational model (Supplementary Fig. 6b) supports this assertion.

We also note that computational model simulation suggests another mechanism underlying selective enhancement. In the computational model, as seen in Supplementary Fig. 6A, the suppression of responses to the surface of the target sphere are dependent on the background input to PV cells which mediate the short-range inter-columnar (inter-receptive field) inhibition (Supplementary Table 3). As this short-range inhibition impinges onto neighboring columns, it is spatially inhomogeneous. For instance, columns responding to the edge will receive short-range inhibition from one side only, whereas columns responding to the center will receive it from all directions. This disparity in lateral inhibition makes column responses to the edge stronger than those to the surface.

The feedback signals from higher visual areas such as V2 and MT (medial temporal visual areas) in primates can also modulate V1 responses^19,20^. V2 reduces V1 responses by enhancing surround suppression^20^, whereas MT enhances V1 responses to moving bars and facilitates figure-ground segregation^19^. That is, V2 and MT regulate V1 responses elicited by moving objects in a similar way VIP cells in V1 do. For instance, the moving objects will elicit stronger responses either when the feedbacks from MT to V1 are stronger or when V1 VIP cell activity is stronger.

Why does the brain use two independent mechanisms to control V1 responses in the same way? Although the feedbacks from MT and VIP cell depolarization lead to higher V1 responses, their influences present different spatial extent. MT may modulate a subset of V1 neurons selectively via cortico-cortical connections, whereas VIP depolarization influences V1 response globally.

When it is necessary to track a specific moving object occupying a subset of visual field, MT, not VIP cells, can enhance V1 responses to it. That is, VIP cells are activated during locomotion but MT may be activated when objects are actually moving inside the visual field. It would be interesting to investigate these two distinct pathways regulating V1 responses to explain the recent observation that V1 neurons respond differently to self-motion and moving objects^21^.

Notably, VIP cells’ depolarization has been also observed in other contextual modulation of sensory cortices. Specifically, VIP cells are nonspecifically activated during conditioning with negative feedbacks^22^, and top-down signals from Cg to V1 target VIP cells mainly^1^, suggesting that VIP cells serve as a unifying mediator for endogenous contextual information originating from other cortices to sensory cortices. However, the exact mechanisms, by which VIP cells contribute to contextual information processing, remain unclear. For instance, SST cells activity increases during Cg activation^1^, whereas it is suppressed during fear conditioning^22^, which remains unexplained.

These two different observations can map onto the high and low intermediate VIP cell depolarization states of the firing rate model. First, in the intermediate VIP cell depolarization condition, SST cells can also be active. As Cg activation depolarizes SST cells as well as VIP cells^1^, the intermediate VIP cell depolarization may cause consistent effects as Cg activation. Indeed, VIP and SST cells may be optimized to promote the competition between them; they mutually inhibit yet promote the identical type to fire more^23^. Second, in the high VIP cell depolarization condition, SST cell activity is uniformly suppressed, which is similar to the observation during fear conditioning^22^. Based on the analyses (Figs. 4 and 5), we propose that sensory cortices may work in two distinct modes. During Cg-activation, sensory neurons become selectively sensitive to some features, which allows V1 neurons to extract behaviorally important information effectively. During fear-conditioning, sensory neurons reliably relay the stimulus inputs, which may help high-order cognitive areas assess the external environments related to the fear conditioning without any biases.

In conclusion, as cognitive functions may depend on interactions among multiple cortical areas^24^, VIP cells’ functional roles could advance our understanding of neural basis of cognitive functions, and we believe that computational models are effective tools to pursue this direction, as we show in this study.

## Methods

### Firing rate model

As seen in Fig. 1A, each population consists of 4 different cell types. For simplicity, we assume all cell types are identical in terms of dynamics of membrane potentials, and their time courses are described by the simple rule.

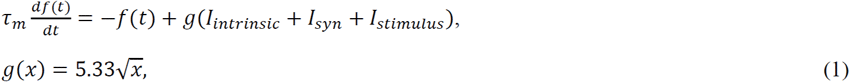

, where *τ*_m_ (the time constant of membrane)=10 ms (the time constant of membrane); where *f* and *g* are the firing rate and gain functions, respectively; where I_intrinsic_, I_sys_ and I_stinulus_ are inputs to the cells. The gain function *g* is obtained by computing the F-I curve of the leaky integrate and the fire neuron implemented by “iaf_psc_exp” included in the peer-reviewed simulator NEST^25^. The square-root is an approximation of the exact analytical solution, but we select this function for two reasons. First, it provides a good approximation as shown in supplementary Fig. 1a and is less computationally intensive than an exact analytical form. Second, it is commonly used as a gain function^11^ I_instrinsic_ is the sum of spiking threshold and background input, which are cell-type specific, as listed in Supplementary Table 2. I_stimulus_ (0.5 pA) is the input representing stimulus presentation, and it is given to Pyr cells only. I_syn_ are synaptic inputs within population and across populations.

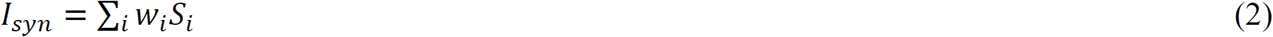

, where *i* runs over all pre-synaptic cells. They are regulated by gating variables *S* and scaled by w_i_. The gating variables *S* evolve according to the activity of presynaptic cell populations^26^, as follows:

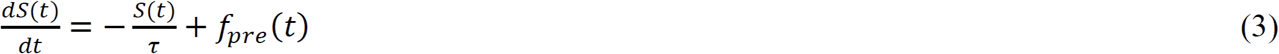

, where *τ* and *f*_pre_ are the decaying time constant and the firing rate of pre-synaptic cells, respectively. The decay time constants are estimated based on physiological data reported in Pfeffer et al. ^9^; this process is discussed elsewhere ^12^. All parameters for synaptic connections are shown in Supplementary Table 1. We solved these equations using the “odeint”, a scipy module included in python.

### Estimates of stimulus-evoked activity

We calculated the stimulus-evoked responses by computing Pyr cell activity during the stimulus period (500-1000 ms). This is normalized in two different ways. First, the signal-to-noise ratio (SNR) is determined by calculating the ratio of population 4 activity to the mean activity of all other populations. Second, the stimulus-period activity is compared to the baseline-period activity by calculating the relative changes in Pyr cell activity (Equation 4):

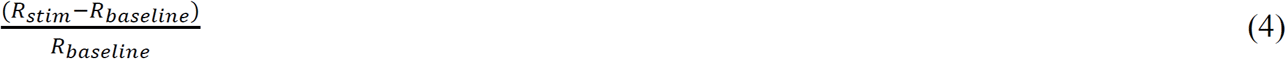

where R_stim_ and R_baseline_ indicate the mean activity of Pyr cells in the stimulus and baseline periods, respectively.

### The spiking neural network model of V1

We extended our earlier V1 model^12^ into 192 column models distributed over 16-by-12 grids by reducing the size of individual columns by a factor of 10 (Fig. 6A). All connections are established randomly^12,15,27^ using the proposed connection probabilities from earlier models^15^. Synaptic strengths used in the model are listed in Supplementary Table 3. The details of cortical column models are discussed elsewhere^12^. Each column receives sensory inputs from 100 thalamic cells, whose firing rate is proportional to the strength of visual inputs within the receptive fields.

For simplicity, we assumed that all thalamic cells are ON cells, and that all thalamic cell populations have non-overlapping receptive fields. Also, thalamic cell populations are distributed over 16-by-12 grids so that they could connect to cortical columns via topographic connections. Each lateral geniculate nucleus (LGN) population consists of 100 thalamic cells, and individual cells induce Poisson spike trains at the fixed rate proportional to the sum of signals (*I*) in the corresponding image patch:

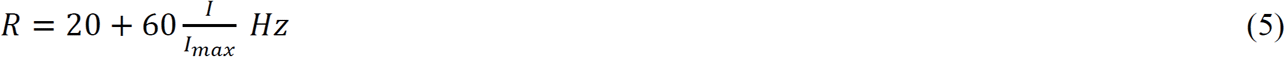

, where *I*_max_ is the maximal value of the sums of intensity of the 192 image patches.

### Visual scene generation

We used POV-Ray to create a simple experimental setup shown in Fig. 6B. The mouse has not been explicitly modelled. Instead, the camera device assumes the role of a mouse’s retina. POV-Ray produces 640-by-480 pixel images in 20 frames during 1 sec in two different conditions, with the width of the image set to 80^o^. The animal translates at constant speed towards the image plane that is perpendicular to the animal’s motion. The five spheres in Fig. 6B are the depicted scene. The center sphere is 50% bigger than all others (Fig. 7A). In both conditions, each frame is 50 ms long and is converted to LGN outputs in 50 ms windows. The size of the receptive field of LGN populations is 40-by-40 pixels of the image so that each frame could be split to 16-by-12 non-overlapping patches.

### Correlations between stimulus inputs and Pyr cell responses

For both firing and computational models, we calculated Pearson’s correlations coefficients between stimulus inputs and Pyr cell responses. In the firing rate model, we record the inputs to and outputs from Pyr cells over time. That is, for each population, the two-time series were collected, from which the correlation was estimated. In the computational model, the correlations were calculated using thalamic outputs 
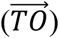
 and column responses 
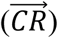
. After recording the column responses depending on 50-ms temporal windows, we converted them into a 1-dimensional vector. Since the center (target) sphere behaves differently from others, we split this 1 dimensional vector into 2 distributions (inside and outside the target sphere). Then, we calculated the correlation coefficients using these two distributions, respectively.

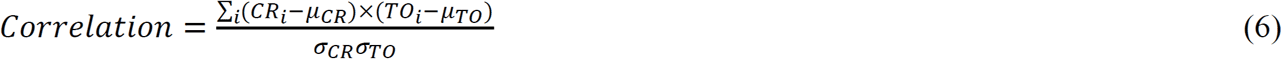

, where *µ* and *σ* are the mean and standard deviation of vector components; where i=pixels inside or outside the center sphere. We instantiated 10 independent networks using the same connectivity, and each network was simulated independently. The correlation was estimated in each simulation.

### Code availability

The simulation codes are available upon request (contact to J. L. at) and will be publicly available in the near future.

## Acknowledgments

We wish to thank the Allen Institute founders, Paul G. Allen and Jody Allen, for their vision, encouragement and support. We would also like to thank Christof Koch for his invaluable feedback on this manuscript.

## Additional Information

### Author Contributions

J.L and S.M. designed the study and wrote the paper. J.L. performed simulations. J.L and S.M. analyzed data.

